# Kinetochore individualization in meiosis I is required for centromeric cohesin removal in meiosis II

**DOI:** 10.1101/2020.07.24.219873

**Authors:** Yulia Gryaznova, Leonor Keating, Sandra A. Touati, Damien Cladière, Warif El Yakoubi, Eulalie Buffin, Katja Wassmann

## Abstract

Partitioning of the genome in meiosis occurs through two highly specialized cell divisions, named meiosis I and II. Step-wise cohesin removal is required for chromosome segregation in meiosis I, and sister chromatid segregation in meiosis II. In meiosis I, mono-oriented sister kinetochores appear as fused together when examined by high resolution confocal microscopy, whereas they are clearly separated in meiosis II, when attachments are bipolar. It has been proposed that bipolar tension applied by the spindle is responsible for the physical separation of sister kinetochores, removal of cohesin protection and chromatid separation in meiosis II. We show here that this is not the case, and initial separation of sister kinetochores occurs already in anaphase I, when attachments are still monopolar, and independently of pericentromeric Sgo2 removal. This kinetochore individualization occurs also independently of spindle forces applied on sister kinetochores, but importantly, depends on cleavage activity of Separase. Crucially, without kinetochore individualization by Separase in meiosis I, oocytes separate bivalents into chromosomes and not sister chromatids in meiosis II, showing that whether centromeric cohesin is removed or not is determined by the kinetochore structure prior to meiosis II.

## Introduction

In meiosis I, chromosomes of different paternal origin (“homologues”) are separated, and in meiosis II, sister chromatids. Because there is no S-phase between the two meiotic divisions, the genome is thus halved and haploid gametes are generated. The meiosis-specific segregation pattern requires the co-orientation of the two sister kinetochores of one homologue to the same spindle pole in meiosis I. Hence, sister kinetochores are attached in a monopolar fashion-this is called mono-orientation; however in meiosis II, sister kinetochores are attached in a bipolar manner, they are bi-oriented (Duro and Marston, 2015; Petronczki et al., 2003). Consequently, the direction of the tension forces applied by the bipolar spindle on sister kinetochore pairs is distinct in meiosis I and II. Sister kinetochores in meiosis I appear fused together when examined by confocal microscopy (Chambon et al., 2013b; Gomez et al., 2007; Lee et al., 2008), whereas they are clearly separated in meiosis II. It was proposed that this separation is due to the application of bipolar tension forces (Gomez et al., 2007; Lee et al., 2008), but it has never been addressed whether sister kinetochores also separate (“individualize”) without bipolar tension, and if they really separate only in meiosis II and not earlier.

To correctly execute both meiotic divisions, the physical connections between chromosomes and sister chromatids are removed in a step-wise manner, from arms in meiosis I and the centromere region in meiosis II. Homologous chromosomes in meiosis I are maintained together by chiasmata, which form on chromosome arms at sites where a meiotic recombination event has taken place and was resolved by a cross-over. Sister chromatids are held together through cohesion, brought about by the cohesin complex containing a kleisin subunit that is cleaved by the protease Separase at metaphase-to-anaphase transition. The separation of chromosomes in meiosis I requires the resolution of chiasmata and the removal of cohesion from chromosome arms because genetic material has been exchanged between chromatids of different parental origin. Importantly, cohesin in the centromere region (where no recombination takes place) has to be preserved, to keep sister chromatids together until anaphase onset in meiosis II (Duro and Marston, 2015; Marston and Wassmann, 2017; Petronczki et al., 2003).

It was shown that protection of centromeric cohesin on homologous chromosome pairs (“bivalents”) in meiosis I depends on recruitment of the phosphatase PP2A-B56 by a Shugoshin (Sgo) family protein, to the centromere region. There, PP2A-B56 counteracts phosphorylation of the meiotic kleisin Rec8, preventing its cleavage by Separase (Clift and Marston, 2011; Keating et al., 2020). As a caveat, it is important to mention that for up to now Rec8 phosphorylation as a requirement for cleavage by Separase *in vivo* has only been demonstrated in yeast and *C. elegans* (Ishiguro et al., 2010; Katis et al., 2010; Rogers et al., 2002; Rumpf et al., 2010). In agreement with this model also applying in higher eukaryotes, inhibition of PP2A-B56 activity, or loss of Sgo2 (which is required for PP2A-B56 recruitment in higher eukaryotes) during the first meiotic division lead to cleavage of Rec8 not only on chromosome arms but also in the centromere region, and hence, precocious sister chromatid separation already in meiosis I (Chang et al., 2011; Lee et al., 2008; Llano et al., 2008; Mailhes et al., 2003; Rattani et al., 2013). Nevertheless, it is still unknown how protection of centromeric cohesin is removed in meiosis II only (a process also called “deprotection” (Chambon et al., 2013b; Jonak et al., 2017; Wassmann, 2013), to allow separation of sister chromatids.

Sgo2 fulfils several interwoven roles in meiosis I, ranging from its contribution to spindle assembly checkpoint (SAC) silencing, delaying tension establishment by the bipolar spindle, Aurora B kinase recruitment, and centromeric cohesin protection (Rattani et al., 2013). Together with the SAC protein Mad2, Sgo2 was shown very recently to additionally function as a Separase inhibitor in mitotic cells, even though it is currently unknown whether this also applies to the meiotic divisions (Hellmuth et al., 2020). Not surprisingly, in oocytes distinct pools of endogenous Sgo2 are recruited to the centromeric region (El Yakoubi et al., 2017), supposedly occupying different roles. We have shown that Sgo2 colocalizing with centromere markers is contributing to bring about cohesin protection in meiosis I. The kinases Mps1 and Bub1 are involved in this recruitment process, and even though both proteins are required for Sgo2 localization, only kinase activity of Mps1 is necessary for implementing its protective role (El Yakoubi et al., 2017).

When analysing the localisation of endogenous Sgo2 in mouse oocytes, we and others have found that Sgo2 is localized to the centromere region of paired sister chromatids (“dyads”) also in meiosis II, where centromeric cohesin has to be cleaved and Sgo2’s protective role is not required (Chambon et al., 2013b; Lee et al., 2008). This is not surprising as such, because apart from preventing Rec8 cleavage, Sgo2’s other roles are most likely also required for correct execution of meiosis II. PP2A-B56 is recruited to centromeres at high levels, too (Chambon et al., 2013b; Lee et al., 2008), suggesting that there must be some kind of mechanism preventing PP2A-B56 action on Rec8, to allow its cleavage by Separase.

Several, not necessarily mutually exclusive models have been proposed for deprotection of centromeric cohesin specifically in meiosis II. First, bipolar tension applied on sister kinetochores in meiosis II, but not meiosis I, was suggested to move Sgo2-PP2A-B56 far enough away from Rec8 to allow its phosphorylation (Gomez et al., 2007; Lee et al., 2008). Second, degradation of Sgo1 and Mps1 at anaphase II onset in an APC/C-dependent manner was proposed to ensure deprotection in budding yeast (Jonak et al., 2017). Third, using a Morpholino-knockdown approach we proposed that I2PP2A/Set, a Histone chaperone and potential PP2A inhibitor, counteracts protection of Rec8 specifically in meiosis II in a tension-independent manner (Chambon et al., 2013b). But apart from the first model, none allows us to understand the key event leading to the deprotection of centromeric cohesin in meiosis II and not meiosis I.

Here, we address the question of which upstream event is required for deprotection of centromeric cohesin in mouse oocytes. We find that bipolar tension is dispensable for step-wise cohesin removal, as deprotection of centromeric cohesin takes also place on tensionless monopolar spindles in meiosis II. We also exclude eviction of Mps1 and pericentromeric Sgo2 as key events mediating deprotection. Crucially, we find that fusion of sister kinetochores is resolved in a Separase-dependent manner already in anaphase I, hence independent of bipolar tension. We find that this kinetochore individualization before entering meiosis II is the key event for allowing centromeric cohesin removal and sister chromatid separation in meiosis II. Rescue experiments of oocytes devoid of Separase (Kudo et al., 2006) demonstrate that Separase activity before entry into meiosis II is required for kinetochore individualization and sister separation in meiosis II. Absence of Separase in meiosis I and its presence only in meiosis II leads to removal of arm cohesin and segregation of bivalents into dyads instead of sister chromatids. In contrast, bivalents with individualized sister kinetochores separate sister chromatids in meiosis II. Hence, the key event for deprotection of centromeric cohesin for sister chromatid segregation in meiosis II takes place already at the final stages of meiosis I in mammalian oocytes. Our data indicate that centromeric cohesin removal is not determined solely by the cell cycle stage (meiosis I or meiosis II), but foremost the chromosome itself.

## Results

### Sister kinetochores individualize already in anaphase I

We wanted to determine whether bipolar tension in meiosis II was indeed required to move Sgo2 and PP2A away from pericentromeric Rec8. Hence, we asked at what time during the transition from meiosis I to meiosis II removal of Sgo2 and PP2A from the pericentromere was observed. When we performed chromosome spreads at the metaphase to anaphase transition of meiosis I, we observed that already in anaphase I sister kinetochores became visible as two separate dots **(Fig. 1A and B)**, even though they are still paired and attached to the same pole on the anaphase I spindle **(Fig. 1B)** (Kitajima et al., 2011). We decided to refer to the first visible separation of the sister kinetochore signals in anaphase I, which was visible both on chromosome spreads and whole mount oocytes, as “kinetochore individualization".

**Figure 1.**
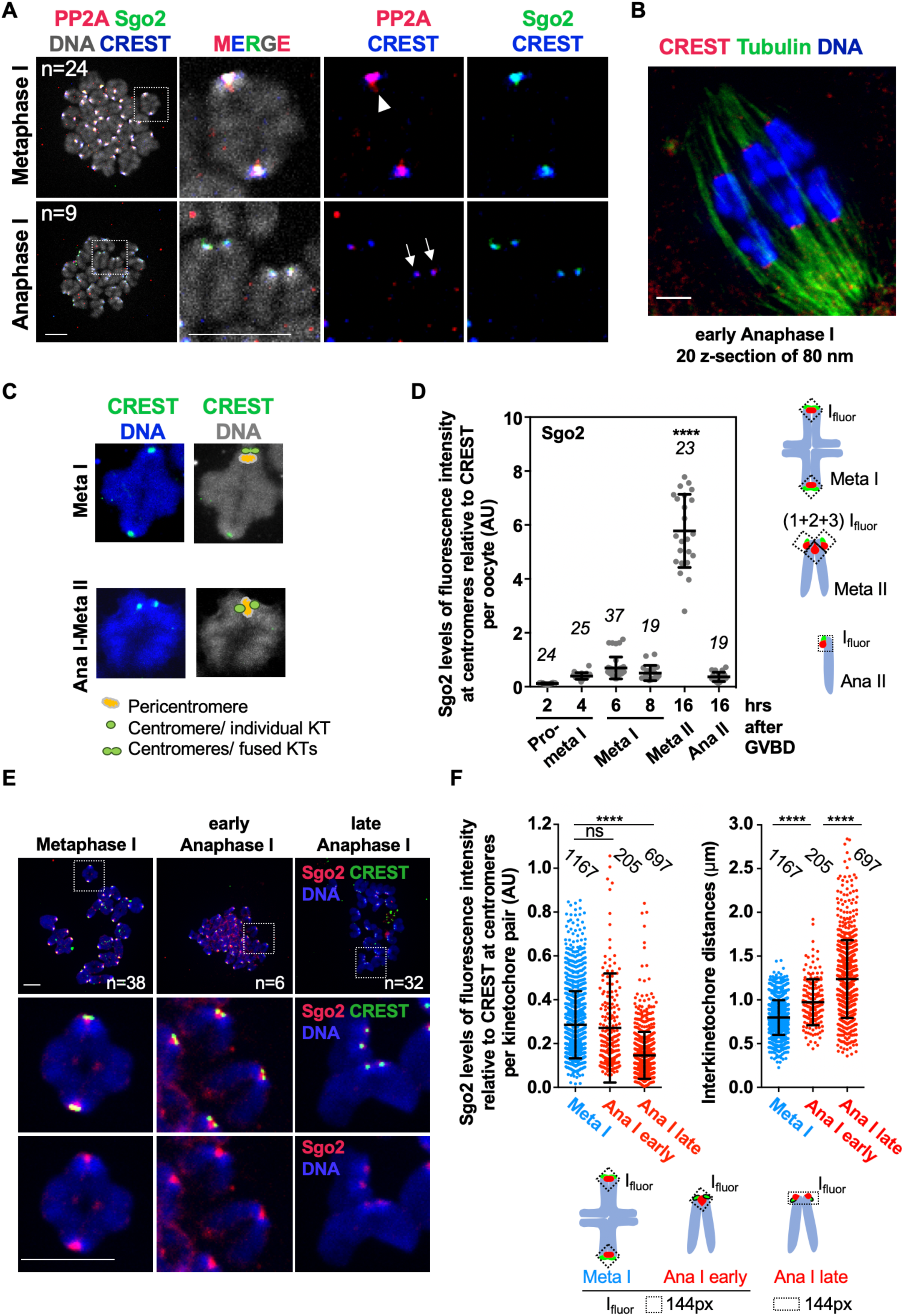
Sister kinetochores individualize already in anaphase I. **A)** Oocytes were fixed for chromosome spreads 8 hours after GVBD, and stained for PP2A-c (red), Sgo2 (green), CREST (blue) and DNA (Hoechst, shown in grey). Spreads were classified into metaphase I and anaphase I, depending on whether chromosome segregation had taken place, or not. Arrowhead indicates fused sister kinetochores, arrows indicate separated sister kinetochores. **B)** Schematic representation of fused or separated sister kinetochores, centromere and pericentromere regions on a bivalent in metaphase I and a dyad in metaphase II. **C)** Example of a whole mount oocyte staining, in early anaphase I after cold-treatment to visualize only cold-stable microtubule fibers, stained with anti-tubulin antibody (green). Kinetochores are stained with CREST serum (red) and chromosomes with Hoechst (blue). Shown is an overlay of 20 z-sections. **D)** Quantification of Sgo2 signals normalized to CREST at centromeres on chromosome spreads from oocytes at the indicated stages of meiotic maturation. Each dot shows mean per oocyte, the number of oocytes analysed is indicated. Below: scheme illustrating how Sgo2 signals were measured at the indicated meiotic stages. I_fluor_ stands for a mean fluorescence intensity of the area within the box (dotted lines, see Material and Methods for quantification approach). **E)** Chromosome spreads as in A), stained for Sgo2 (red), CREST (green), and DNA (Hoechst, in blue). Spreads in early anaphase I and late anaphase I were classified, depending on whether dyads of two chromosome sets had been found close to each other as one group, or were scattered apart into two groups. In early anaphase I, Sgo2 is found in between sister kinetochores that are already separated. **F)** Quantification of D) showing Sgo2 signal relative to CREST per kinetochore pair (dot plot on the left), and sister kinetochore distance (dot plot on the right) at the indicated stages of meiosis I. The number of kinetochore pairs analysed is indicated. On the right: scheme of Sgo2 signal measurements. On each graph mean is shown, error bars are ± SD, asterisks indicate significant difference (p<0,0001) and ns - not significant, according to t-test. Scale bars, 10μm.

This separation of sister kinetochores has been observed before in metaphase II oocytes (Chambon et al., 2013b; Kim et al., 2015; Lee et al., 2008) and was thought to be due to the bipolar tension applied on dyads as they align at the metaphase plate and are oriented towards the opposite poles of the bipolar spindle in meiosis II (Lee et al., 2008). The cohesin protector Sgo2 and PP2A co-localized with CREST, which marks proteins at the centromere and inner kinetochore in metaphase I. Here we show that both Sgo2 and PP2A were removed from the pericentromere in anaphase I **(Fig. 1A and C)**. Altogether, our result was unexpected because 1) sister kinetochores separated already on the anaphase I spindle, when they are attached to the same pole **(Fig 1B)**, contradicting the hypothesis that bipolar tension is required for sister kinetochore separation, 2) centromeric cohesin between sister kinetochores has to be protected in anaphase I, when Separase should still be active, given that Cyclin B1 and Securin are absent as oocytes exit meiosis I. The removal of Sgo2 from this region -the pericentromere **(Fig. 1C)** - in anaphase I already, came therefore as a surprise.

To better understand how this kinetochore individualization and Sgo2 localization are related, we first re-analysed the endogenous centromeric localization of Sgo2, in synchronized mouse oocytes cultured *in vitro* and fixed at key stages of meiosis I and II. Prophase I arrested oocytes were released to enter meiosis I and allowed to progress into meiosis II, until cell cycle arrest in metaphase II (CSF arrest), where they await fertilization. Examination of endogenous Sgo2 localization on chromosome spreads showed that Sgo2 is recruited to the centromere region throughout meiosis I, as shown before **(Fig. 1D)** (El Yakoubi et al., 2017; Lee et al., 2008; Lister et al., 2010; Rattani et al., 2013). We prepared chromosome spreads at the metaphase to anaphase transition of meiosis I, and classified spreads into metaphase I, and early and late anaphase I, depending on whether separating dyads were found right next to each other, or dyads had moved away from each other into two independent pools. Sgo2 at the centromere can be detected throughout anaphase I, whereas Sgo2 at the pericentromere was present in early anaphase, and disappeared in late anaphase **(Fig. 1E)**. Kinetochore individualization started in early anaphase I, and at this stage, sister kinetochores that have separated, but still harbour Sgo2 in the pericentromere, were detected **(Fig. 1F and G)**. Hence pericentromeric Sgo2 is removed too late for being the initial trigger for sister kinetochore individualization in meiosis I.

### Localization of Sgo2 at the pericentromere is dynamic

Strikingly, staining of endogenous Sgo2 showed an approximately 5-fold increase of Sgo2 levels in metaphase II, compared to metaphase I **(Fig. 1D)**. The pericentromeric fraction of Sgo2 that had been removed in anaphase I, was re-established in metaphase II **(Fig. 2A)**. Pericentromeric Sgo2 removal in metaphase II may be essential for cohesin deprotection in anaphase II, therefore we further analysed endogenous Sgo2 localization in meiosis II. Anaphase II takes place as soon as fertilization occurs, or alternatively, after chemical activation of oocytes cultured *in vitro*. Sgo2 protein levels decreased again in anaphase II **(Fig. 1D)**, but importantly, Sgo2 was nevertheless still present in the centromere region, colocalizing with the CREST signal **(Fig. 2A)**. The fraction of Sgo2 that disappeared was indeed the one found in between the two kinetochore signals of the sister chromatids, at the pericentromere **(Fig. 1C and 2A)**. Our data indicate that there are two pools of Sgo2 as oocytes progress from meiosis I into meiosis II: one that remains stably associated with centromeres, and another one at the pericentromere that is highly dynamic, removed when oocytes exit meiosis I, and relocated when oocytes enter meiosis II, to be removed again in anaphase II. Hence, pericentromeric but not centromeric Sgo2 removal may be required for deprotection of cohesin in meiosis II.

**Figure 2.**
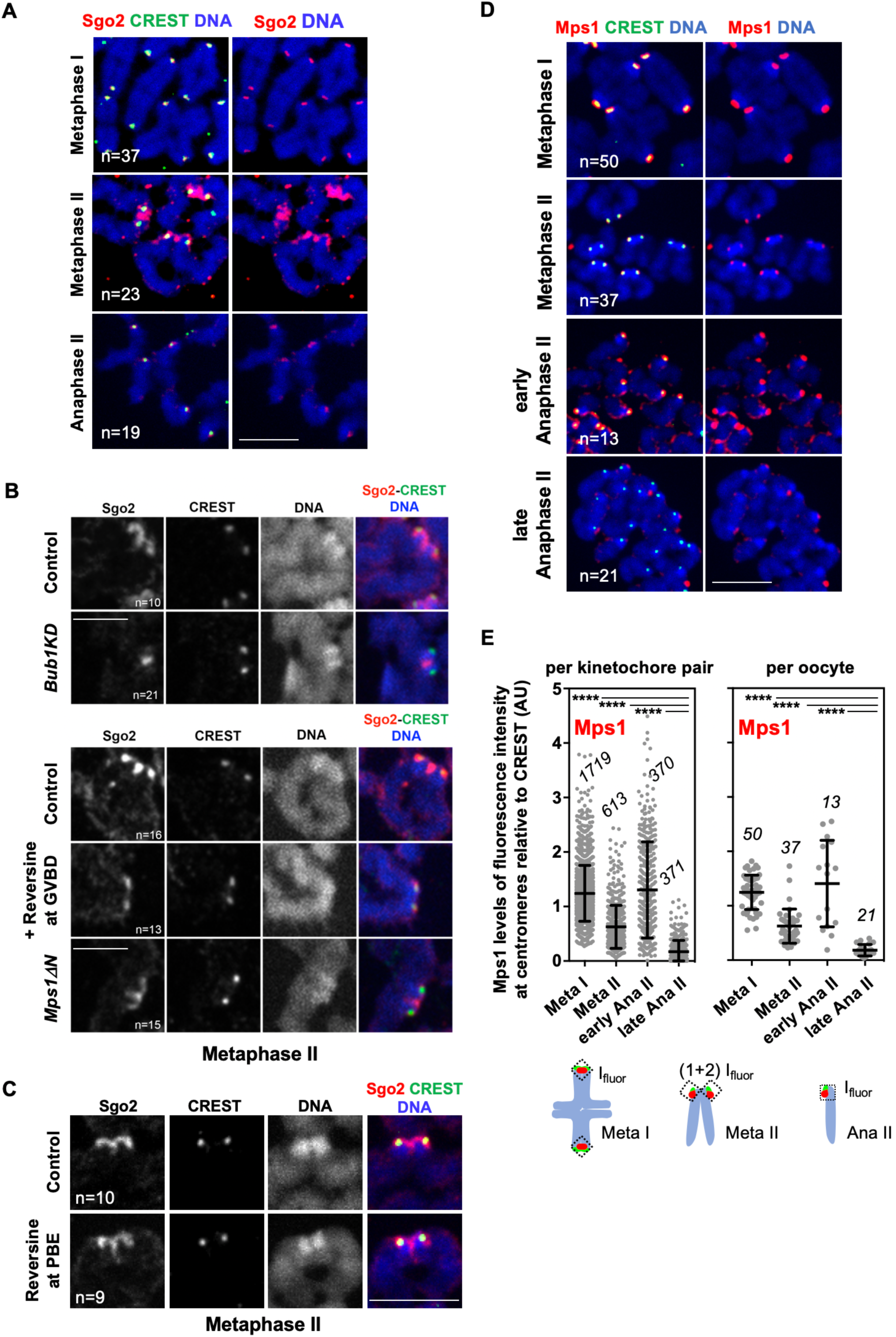
Pericentromeric Sgo2 localized by Mps1, is dispensable for sister chromatid segregation. **A)** Chromosome spreads at the indicated stages of meiosis, stained for endogenous Sgo2 (red), centromeres with CREST antibody (green) and DNA with Hoechst (blue). **B)** Bivalents of metaphase II chromosome spreads of control, Bub1KD, Mps1ΔN and Reversine-treated oocytes (from GVBD onwards) stained with Sgo2 antibody (red), CREST antibody (green) and Hoechst (blue). **C)** Same as B), except that oocytes were treated with Reversine from GVBD + 8 hours onwards, corresponding to the time of anaphase I onset. PBE: Polar Body Extrusion. **D)** Chromosome spreads at the indicated stages of meiosis, stained for endogenous Mps1 (red), centromeres with CREST (green) and DNA with Hoechst (blue). For early and late anaphase II spreads oocytes were chemically activated and fixed 25min or 1h later respectively. **E)** Quantification of C) showing Mps1 signal relative to CREST per kinetochore pair (per single kinetochore in anaphase II) (left dot plot), and Mps1 signal averaged per oocyte (right dot plot) at the indicated time points. The number of kinetochore pairs (kinetochores in anaphase II) and oocytes analysed is indicated. The scheme of measurements is shown below. On each graph mean is indicated, error bars are ± SD, **** is significant difference (p<0,0001) according to t-test. Scale bars, 10μm. For spreads, n indicates number of spreads analysed.

### Sgo2 localized by Mps1 kinase to the pericentromere is removed at anaphase II onset

In meiosis I, different pools of endogenous Sgo2 can be distinguished in the centromere region. We have shown previously that Sgo2 brought to the centromere in a Mps1 kinase-dependent manner is required for centromeric cohesin protection in meiosis I (El Yakoubi et al., 2017). There is another pool of Sgo2 localized in a Bub1 kinase-dependent manner, but this one is not required for cohesin protection in meiosis I (El Yakoubi et al., 2017). It was attractive to speculate that the pool of Sgo2 that is removed in meiosis II is the one localized there by Mps1, hence potentially protecting centromeric cohesin until anaphase II onset. To address this issue we asked whether also in metaphase II, distinct pools of Sgo2 are localized by Mps1 and Bub1 kinase activities, such as in meiosis I. To address this issue we made use of two mouse models, one harbouring a kinase-dead version of Bub1 instead of wild type Bub1 (El Yakoubi et al., 2017; Ricke et al., 2012), and the other one expressing only a truncated version of Mps1 (Mps1ΔN) that cannot localize to kinetochores, but still harbours kinase activity (Hached et al., 2011). Additionally, we added the Mps1 inhibitor Reversine to wild type oocytes during the entire meiotic maturation from GVBD onwards, to inhibit Mps1 kinase activity, but not its localization. Indeed, similar to meiosis I (El Yakoubi et al., 2017), Bub1 and Mps1 kinases preferentially localize distinct pools of Sgo2, but this time to the centromere and pericentromere, respectively **(Fig. 2B, Fig. S1)**. Hence, the fraction of Sgo2 that is removed in anaphase II corresponds to Sgo2 localized there by Mps1 kinase activity, suggesting that this pool, in analogy to meiosis I, may confer protection until anaphase II onset. Kinetochore localization of Mps1 is not specifically required for localizing Sgo2 to the pericentromere, as Mps1ΔN oocytes do not preferentially lose Sgo2 from the pericentromere. Mps1ΔN oocytes do not show loss of centromeric cohesin protection in meiosis I or II (El Yakoubi et al., 2017), further indicating that kinetochore localization of Mps1 is not required for protection. Together, our data indicate that the Mps1-kinase activity dependent fraction of Sgo2 at the pericentromere, and not the one at the centromere, is removed for sister separation in meiosis II.

We were wondering at what stage of meiotic maturation Mps1 kinase activity was necessary to obtain Sgo2 at the pericentromere in meiosis II. When we inhibited Mps1 only during meiosis II by adding Reversine at the time of meiosis I Polar Body extrusion, we did not lose the pericentromeric fraction of Sgo2 **(Fig. 2C)**, unlike what we observed upon inhibition of Mps1 in meiosis I already **(Fig. 2B)**. This indicates that modifications brought about by Mps1 in meiosis I and not meiosis II are required for proper pericentromeric localization of Sgo2 in metaphase II. As kinetochore localization of Mps1 was not required **(Fig. 2B)**, and taking into account the increase of Sgo2 levels in the centromere region in meiosis II **(Fig. 1D)**, we conclude that cytoplasmic activity of Mps1 in meiosis I but not meiosis II is required for recruiting Sgo2 from the cytoplasm to the pericentromere.

Even though kinetochore localization of Mps1 is not required for Sgo2 localization at the pericentromere, inhibition and/or removal of Mps1 may still be necessary for removal of pericentromeric Sgo2 in meiosis II. Indeed, in *S. cerevisiae*, degradation of the protector Sgo1 and Mps1 at anaphase II onset was proposed to contribute to deprotection of centromeric cohesin (Jonak et al., 2017). However, throughout metaphase II-to-anaphase II transition Mps1 remains on kinetochores of separating sister chromatids in mouse oocytes, and was never found at the pericentromere. Mps1 kinetochore localization was detected at high levels at the very onset of anaphase II during a very short time window and, as oocytes progressed through anaphase II, Mps1 disappeared quickly, resulting in some kinetochores that had already lost their Mps1 staining and others still harbouring Mps1 **(Fig. 2D and E)**. Altogether we think it is unlikely that deprotection of centromeric cohesin in oocytes is brought about by degradation of Mps1, because sister chromatids separate before Mps1 disappears. Again, this is in agreement with our previous data that kinetochore localization of Mps1 is not required to prevent precocious sister chromatid separation (El Yakoubi et al., 2017).

### Bipolar tension is dispensible for step-wise cohesin removal

It was proposed that deprotection of centromeric cohesin is brought about by bipolar tension applied by the spindle in meiosis II, to physically move Sgo2 and PP2A away from pericentromeric Rec8, hence allowing Rec8 phosphorylation and its conversion into a cleavage substrate for Separase (Gomez et al., 2007; Lee et al., 2008). This does not fit with our data, where Sgo2 and PP2A are removed from the pericentromere already in anaphase I, without bipolar tension **(Fig. 1A)**, and come back in the presence of bipolar tension (**Fig. 2A**)(Chambon et al., 2013b). To clarify this point, we set out to test whether indeed the way tension forces are applied in meiosis I or II determines if the meiotic cohesin subunit Rec8 is removed from chromosome arms or from the centromere region.

First, to determine whether attachment of chromosomes in meiosis I to both poles of the bipolar spindle is a prerequisite for arm cohesin removal, we treated prometaphase I mouse oocytes with the Eg5 inhibitor STLC to induce monopolar spindles, where bivalents with fused sister kinetochores were attached on monopolar spindles (Vallot et al., 2018). Individual kinetochore fibers forming end-on attachments were clearly visible by high resolution confocal microscopy of cold-treated spindles **(Fig. 3A)**. Missing tension prevents timely anaphase I onset, because Aurora B-dependent error correction leads to Spindle Assembly Checkpoint (SAC) activation (Vallot et al., 2018), therefore we added Reversine in metaphase I to override the SAC and allow anaphase I onset in oocytes harbouring monopolar spindles. Short Reversine treatment just before anaphase I onset does not interfere with cohesin protection in meiosis I (El Yakoubi et al., 2017). Upon SAC override, bivalents separated into dyads on monopolar spindles, and arm cohesin was removed, as visualized by Rec8 staining **(Fig. 3B and C)**. When we examined bivalents that separated into dyads on a monopolar spindle, we observed that sister kinetochores that appeared fused together, were clearly separated once arm cohesin was gone **(Fig. 3C)**, further confirming that sister kinetochore individualization does not require bipolar attachment. Rec8 was not removed from the centromere region holding sister chromatids together, and no sister chromatid segregation was observed **(Fig. 3C)**.

**Figure 3.**
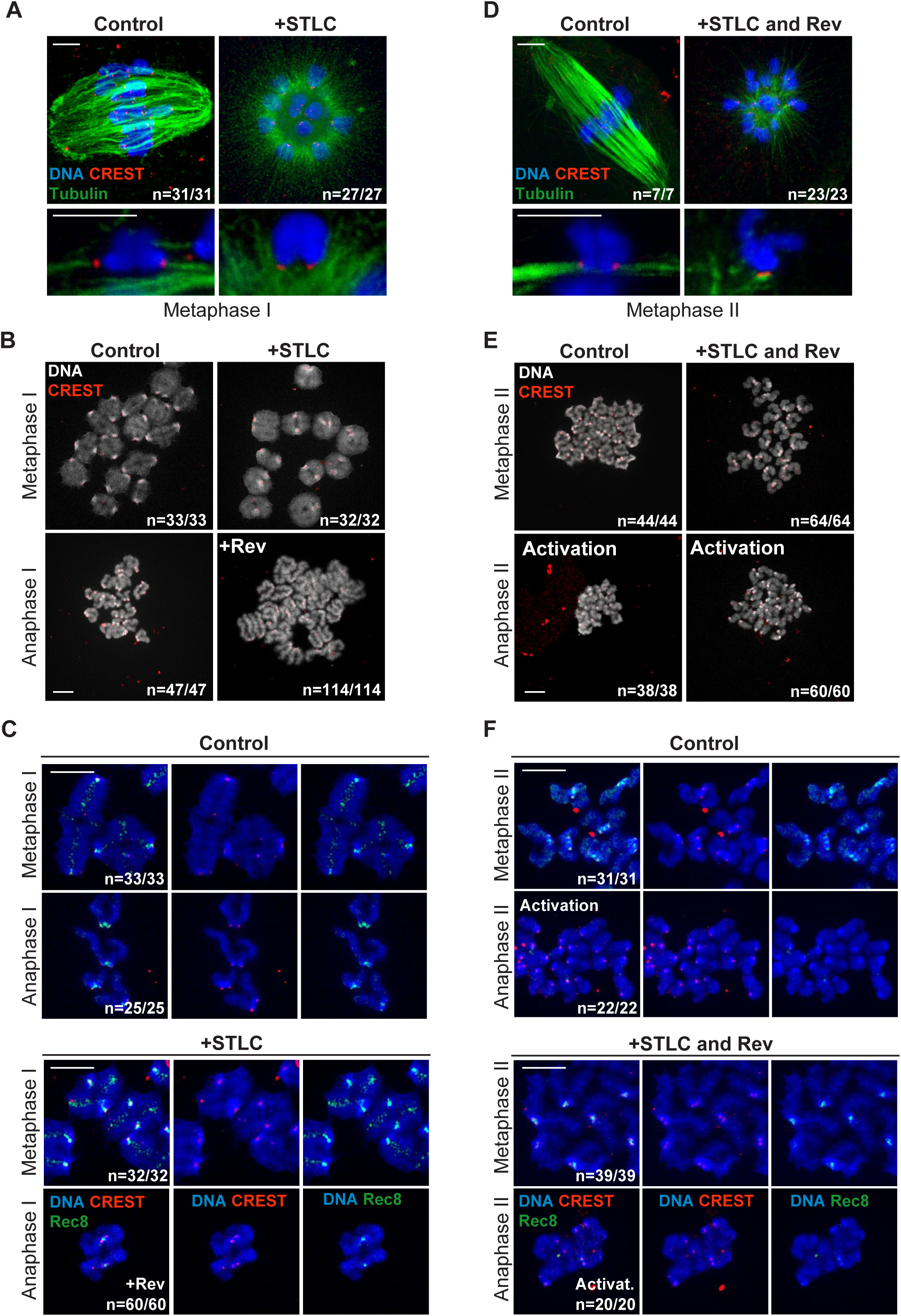
Step-wise cohesin removal occurs independently of tension forces applied by a bipolar spindle. **A and D)** Oocytes in meiosis I (6 hours after GVBD) or meiosis II (left and right panels, respectively) were treated with STLC for 2.5 hours and fixed for whole mount immunofluorescence. Cold stable microtubules were stained with an antibody against tubulin (green), centromeres with CREST (red) and DNA with Hoechst (blue). Scale bar indicates 5 µm. **B and E)** Chromosome spreads at metaphase and anaphase of meiosis I (6 and 9 hours after GVBD) or meiosis II (left and right panels, respectively), in the presence or absence of STLC (for 2.5 hours) and Reversine (+Rev., for 1 hour) as indicated. Spreads were stained with CREST (red) and Hoechst (grey). Single chromosomes are present in anaphase I and single sister chromatids in anaphase II. **C and F)** Chromosome spreads exactly as described in B and E), except that spreads were additionally stained with an antibody against Rec8 (green). The CREST signal appears in red and DNA in blue. Scale bar indicates 10 µm. In all panels the number of oocytes analysed and showing the same phenotype is indicated.

To address whether application of bipolar tension on dyads in meiosis II is required for centromeric cohesin removal we performed the same experiment as above, but in meiosis II: monopolar spindles were obtained through STLC treatment of metaphase II arrested oocytes, SAC response was inhibited through Reversine treatment, and oocytes were activated to undergo the second meiotic division. Microtubule attachments to the monopolar spindle were verified by high resolution confocal microscopy of cold-stable microtubule fibers **(Fig. 3D)**. Remarkably, our experiment showed that sister chromatid segregation and hence, centromeric cohesin removal, does not require bipolar attachment of dyads **(Fig. 3E)**. Centromeric Rec8 was undetectable upon activation, independently of spindles being mono-or bipolar **(Fig. 3F)**. Accordingly, sister chromatids came apart on a monopolar spindle, contradicting the hypothesis that tension-dependent removal of cohesin protection through bipolar attachment is required for centromeric cohesin cleavage, at least in mouse oocytes.

### Individualization of sister kinetochores depends on Separase

Our data indicate that neither removal of pericentromeric Sgo2, inactivation of Mps1, nor bipolar tension are essential for step-wise cohesin removal in oocytes. We hypothesized that kinetochore individualization in meiosis I is determining whether centromeric cohesin can be removed in the following division. It was attractive to speculate that Separase activity is required for individualizing sister kinetochores in anaphase I, and that this is the key event to allow centromeric cohesin removal in meiosis II. To address this hypothesis, we first asked whether oocytes without Separase contained fused or individualized kinetochores, using mice harbouring an oocyte-specific invalidation of Separase (Sep-/-). Oocytes without Separase progress into meiosis II, because loss of Separase does not prevent progression through meiosis I into meiosis II, even though chromosome segregation does not take place. Cyclin B1 and Securin accumulate as oocytes progress into metaphase I, are degraded and re-accumulate in control and Sep-/-oocytes with similar kinetics **(Fig. S2)** (Kudo et al., 2006). Furthermore, upon hormonal stimulation of female mice to induce meiotic maturation of oocytes *in vivo*, control and Sep-/-oocytes move from ovaries into the oviduct and are found as cumulus-enclosed oocytes awaiting fertilization **(Fig. 4A)**. In Sep-/-oocytes, bivalents were still attached in a monopolar manner and sister kinetochores of bivalents remained fused together even though oocytes were in metaphase II **(Fig. 4B)**. This result indicates that Separase is required for sister kinetochore individualization prior to metaphase II in oocytes matured *in vivo*.

**Figure 4.**
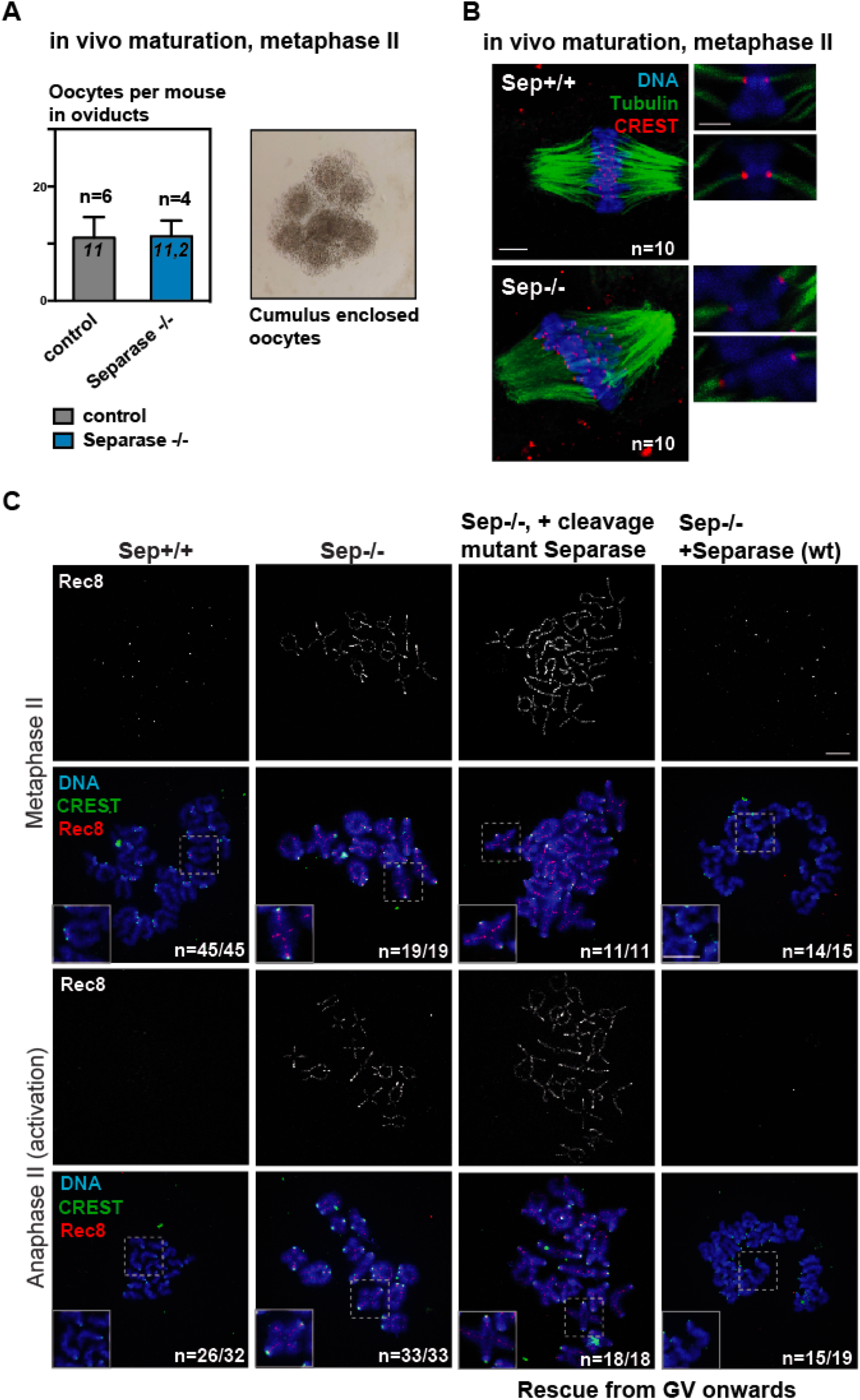
Oocytes without Separase can be activated to undergo meiosis II and separate sister chromatids upon re-expressing of wild type Separase from GV onwards. **A)** Metaphase II oocytes from mice carrying a conditional, oocyte-specific invalidation of Separase (Separase ^flox/flox^ Zp3 Cre^+^, or Sep-/-) and oocytes from litter mates (Separase ^flox/flox^, or Sep+/+) were obtained after superovulation. *In vivo* matured Sep-/-oocytes are found in oviducts at comparable numbers, enclosed by cumulus cells. **B)** Whole mount immunofluorescence to stain cold stable microtubule fibers after *in vivo* maturation. Microtubules were stained with tubulin (green), centromeres with CREST (red) and DNA with Hoechst (blue). Bivalents are attached in a monopolar fashion. Scale bars indicate 5 μm. **C)** Metaphase-to-anaphase transition of meiosis II in oocytes of the indicated genotype cultured *in vitro* and injected with the Separase constructs indicated in GV. Chromosome spreads are stained with CREST antiserum (green), and Rec8 antibody (red). DNA was stained with DAPI (blue). The number of oocytes analysed and the proportion of oocytes with the same phenotype is indicated. Scale bars indicate 10 µm.

Not surprisingly, in the absence of Separase activity, oocytes were unable to segregate chromosomes or sister chromatids upon activation **(Fig. S3 and 4C)**. Meiosis I chromosome segregation, kinetochore individualization, and meiosis II sister chromatid segregation were rescued when Separase was re-expressed from GV onwards, but not when a catalytically inactive Separase mutant was used, showing that Separase is essential to remove centromeric cohesin also in meiosis II **(Fig. S3 and 4C)**. Our data additionally indicate that Separase is required for kinetochore individualization in meiosis I, *in vitro*.

### Bivalents with fused kinetochores separate into dyads, not sister chromatids in meiosis II

We set out to address whether Separase activity during the first meiotic division -supposedly to bring about kinetochore individualization-is required for deprotection and separation of sister chromatids in meiosis II. For this, we asked how Separase-/-oocytes segregate bivalents when rescued with exogenously expressed Separase only in meiosis II, by injecting metaphase II arrested oocytes with mRNA coding for wild type Separase. Injected metaphase II arrested Separase-/- oocytes were activated to undergo anaphase II and then examined by chromosome spreads to address whether chromosomes or sister chromatids were separated **(Fig. S3)**. Remarkably, under these conditions chromosomes were separated and Rec8 was removed from chromosome arms, whereas sister chromatids remained paired with centromeric Rec8 that was not removed by Separase **(Fig. 5A)**. Hence, if Separase is absent in meiosis I, it removes arm cohesin instead of centromeric cohesin in meiosis II.

**Figure 5.**
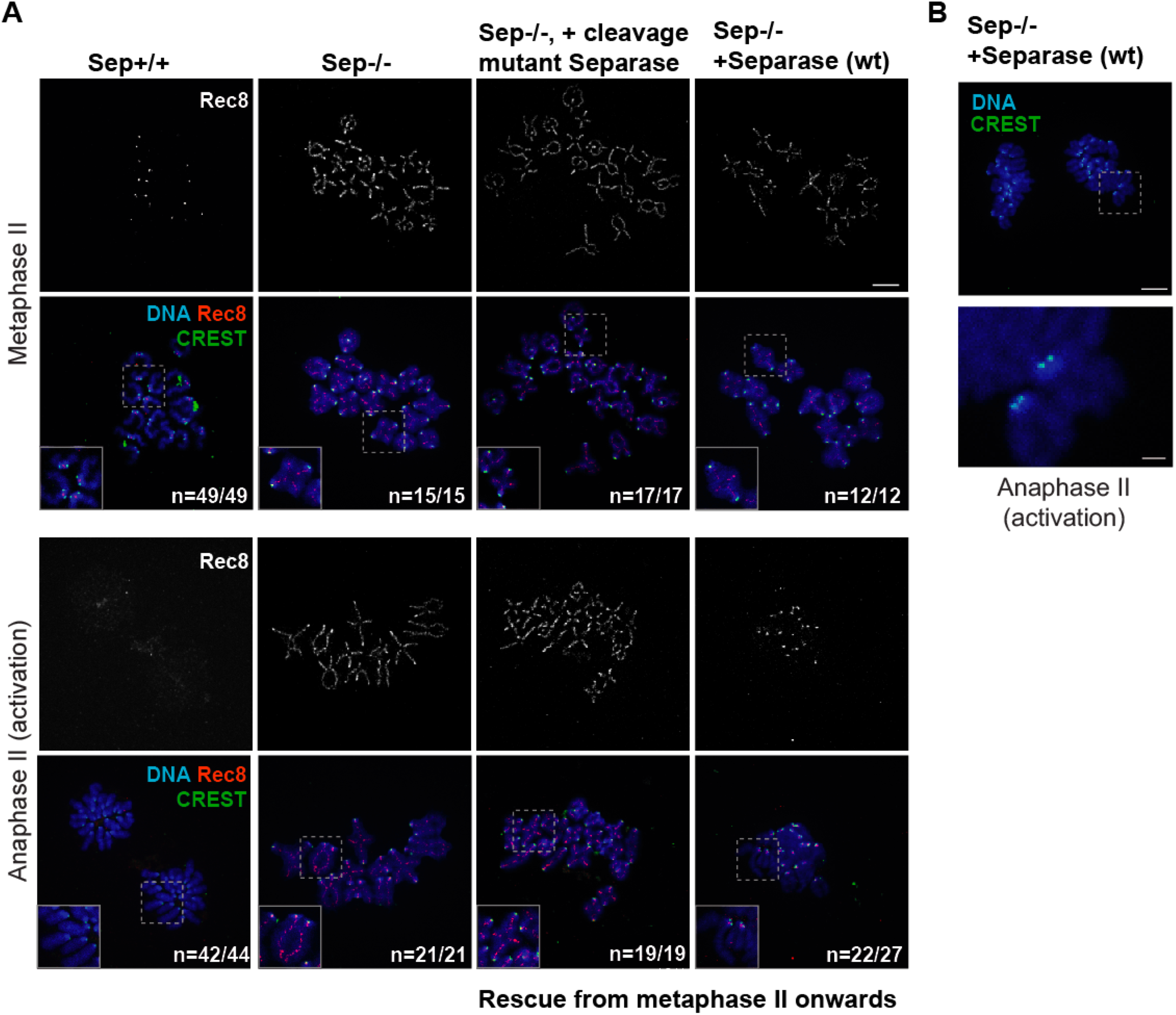
Separase activity during meiosis I is required for centromeric cohesin removal in meiosis II. **A)** Sep+/+ or Sep-/- oocytes were matured *in vitro* until metaphase II. Sep-/- oocytes were injected with wild type Separase or cleavage mutant Separase encoding mRNAs in metaphase II. Oocytes were either fixed in metaphase II, or activated and fixed in anaphase II. Spreads were stained with antibodies against Rec8 (red), CREST serum (green), and DAPI (blue) to label DNA. The number of oocytes analysed and the proportion of oocytes with the same phenotype is indicated. Scale bars indicate 10 µm. **B)** Representative image of kinetochore individualization in Sep-/- oocytes injected with wild type Separase in metaphase II and fixed after activation in A). Shown is staining with CREST serum (green), and DAPI (blue) to label DNA. Scale bars indicate 10μm on the top panel and 2 μm on the bottom panel.

### Sister kinetochore individualization can also occur in meiosis II

Closer examination of sister kinetochores showed that arm cohesin removal and separation of chromosomes in meiosis II in Separase-/- oocytes rescued from metaphase II onwardds also led to sister kinetochore individualization, such as usually observed in meiosis I **(Fig. 5A and B)**. Injecting catalytically inactive Separase did not lead to kinetochore individualization upon activation **(Fig. 5A)**. We conclude that in addition to arm cohesin removal, sister kinetochore individualization requires Separase activity in meiosis I. Both events can occur in meiosis II, if Separase is absent in meiosis I, but our data indicate that Separase cannot induce sister kinetochore individualization and remove centromeric cohesin at the same time.

### Bivalents with individualized kinetochores separate into sister chromatids in meiosis II

We propose that Separase activity at anaphase I is required for kinetochore individualization, and this individualization is a prerequisite for deprotection of centromeric cohesin in meiosis II. In this case, oocytes harbouring bivalents in metaphase II with individualized kinetochores such as usually the case for dyads, should not segregate chromosomes, but sister chromatids.

Overexpression of mitotic Sgo1 was shown to interfere with arm cohesin removal in mouse oocytes in a previous study, by recruiting PP2A to chromosome arms, hence resulting in the presence of bivalents in meiosis II (Xu et al., 2009). In oocytes, Sgo2 and not Sgo1 is essential for protecting centromeric cohesin (Llano et al., 2008), hence we refrained from using Sgo1 to avoid creating artefacts and asked whether overexpression of meiotic Sgo2 equally allowed us to obtain bivalents in meiosis II. Our data above indicated that Sgo2 does not regulate sister kinetochore individualization, as individualization of kinetochores in meiosis I was not due to removal of Sgo2 from the pericentromere **(Fig. 1E and F)**. Consequently, overexpression of Sgo2 was expected to lead to failures in separating chromosomes, yet allow individualization of sister kinetochores. To address this issue we cultured oocytes overexpressing GFP-Sgo2 from GV onwards. GFP-Sgo2 localized to whole chromosomes in prometaphase and metaphase I, as visualized by following the first meiotic division by live-imaging **(Fig. 6A)**, similar to what has been observed before (Rattani et al., 2017; Rattani et al., 2013). Indeed GFP-Sgo2 overexpression prohibited complete cohesin removal from chromosome arms **(Fig. 6B)**. As a result, some bivalents that were not able to separate in meiosis I, were present in metaphase II in roughly half of the injected oocytes **(Fig. 6C)**. Importantly though, passing through meiosis I induced the individualization of sister kinetochores on all chromosomes, including the bivalents that did not separate **(Fig. 6C)**. This confirms that indeed, individualization of kinetochores in meiosis I was not affected by Sgo2 overexpression.

**Figure 6.**
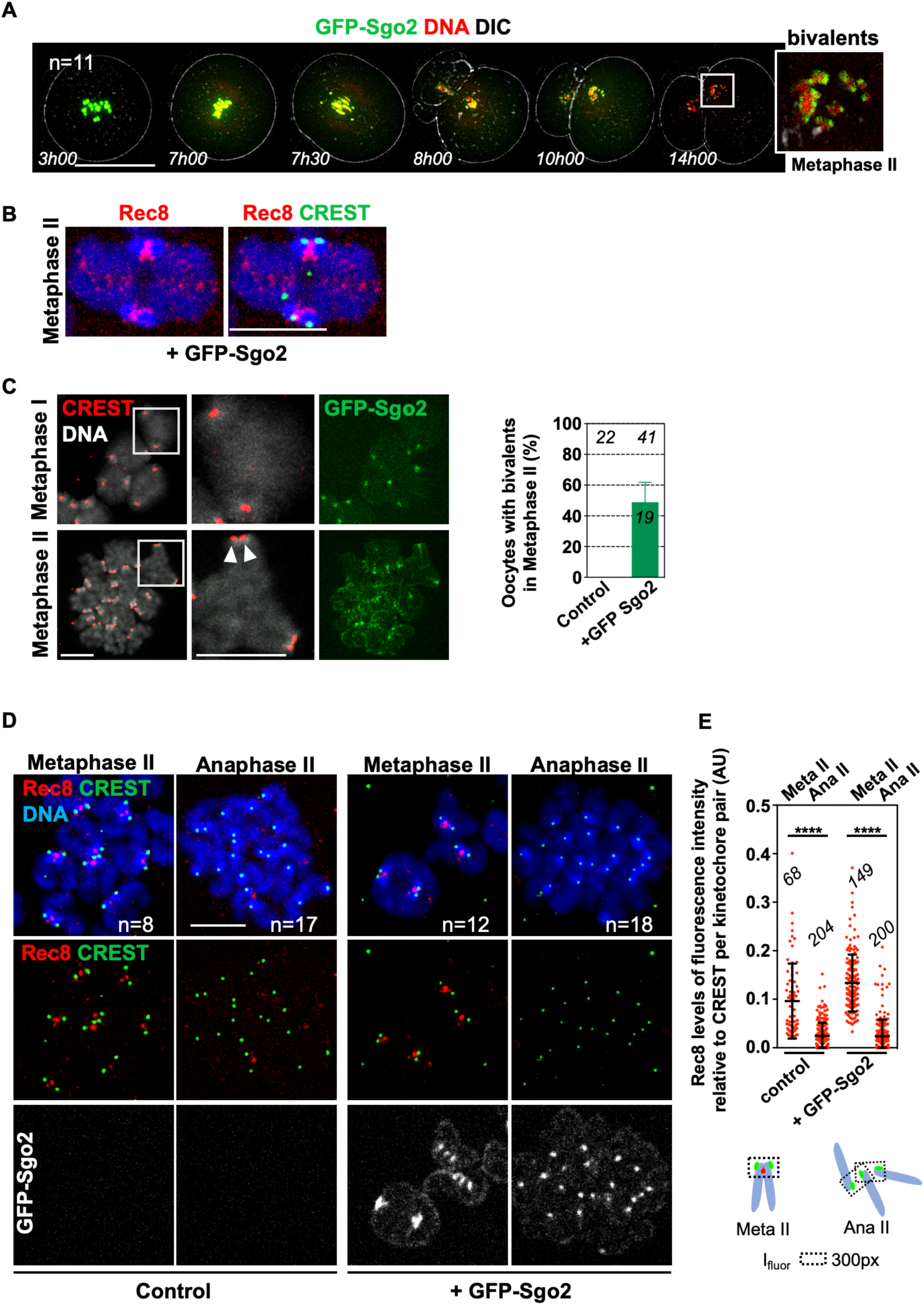
Centromeric cohesin is not protected on bivalents with individualized kinetochores in meiosis II. **A)** Live imaging movie of GFP-Sgo2 expressing oocytes (from GV onwards) undergoing meiosis I. Chromosome movements were followed by the probe for DNA, SirDNA in far red, GFP-Sgo2 appears in green. Shown are overlays of DNA, GFP and DIC channels, each timepoint comprised of 11 z-sections of 3μm and 1 z-section in DIC. Timepoints were taken every 10min, shown are selected timepoints such as indicated. **B)** Representative images showing Rec8 localization on a bivalent chromosome in metaphase II of a GFP-Sgo2 expressing oocyte. Spreads were stained with Rec8 antibody (red), CREST antibody for centromeres (green), Hoechst for DNA (blue). **C)** Early metaphase I (4hrs after GVBD) and II spreads after GFP-Sgo2 expression (from GV onwards). Spreads were stained with CREST antibody (red), Hoechst for DNA (grey), GFP-Sgo2 was visualized by GFP fluorescence (green). Arrowheads mark sister kinetochore separation. The graph below shows the percentage of metaphase II spreads containing bivalent chromosomes. The number of control and GFP- Sgo2 expressing oocytes analysed is indicated. Error bar is +SD. **D)** Chromosome spreads of oocytes expressing GFP-Sgo2 (from GV onwards) in metaphase II, and upon activation. Spreads were stained with antibody against Rec8 (red), CREST antibody (green), and Hoechst (blue). **E)** Quantification of Rec8 signal found around and in between CREST signals that were close to each other. The number of kinetochore pairs analysed is indicated, as well as mean ± SD, **** (p<0,0001) according to Student’s t-test. Scale bars: A) 100μm, B) C) D) 10μm.

Upon activation, dyad separation in meiosis II was not perturbed by GFP-Sgo2 overexpression, even though GFP-Sgo2 was able to localize to the pericentromere in anaphase II **(Fig. 6D)**. Thus, if centromeric cohesin were still protected in meiosis II in a Sgo2-dependent manner, the underlying molecular mechanism must be more complex than mere localization of Sgo2 to the pericentromere. Crucially though, upon activation, bivalents separated into sister chromatids, because no dyads were observed in anaphase II spreads upon activation. All detectable Rec8, on arms and in the centromere region, was removed **(Fig. 6D and E)**. In conclusion, individualization of sister kinetochores in meiosis I leads to centromeric cohesin removal in meiosis II, even on bivalents.

## Discussion

Our study was motivated by a quest to understand how centromeric cohesin is removed only in meiosis II in mammalian oocytes. None of the current models provided a satisfying answer to this question yet. It was proposed that protection of centromeric cohesin is mediated by recruitment of Sgo2/PP2A to the fraction of Rec8 that is protected, and centromeric cohesin is deprotected by removal of Sgo2/PP2A in meiosis II. Bipolar tension was proposed to move Sgo2 together with PP2A away from centromeric Rec8 in both male and female meiosis II (Gomez et al., 2007; Lee et al., 2008). This model was the only one explaining how centromeric cohesin would be deprotected in meiosis II, but not meiosis I. However, this model does not fit with observations in mouse oocytes, where Sgo2 and PP2A co-localize with Rec8 at the pericentromere in meiosis II (Wassmann, 2013). We show here that Sgo2 and Mps1 remain localized to the centromere throughout anaphase II, and anaphase II onset can take place without removing Mps1. This makes it rather unlikely that Sgo2 and Mps1 would just be degraded for deprotection in oocytes. We can speculate that the multiple roles Sgo2 occupies in mammalian oocytes (Rattani et al., 2013) require another layer of regulation. For example, Sgo2 at the pericentromere may be subject to post-translational modifications regulating its capacity to protect centromeric cohesin, and these modifications would need to be removed for deprotection.

Based on our observation that sister kinetochores come apart already in anaphase I, when they are still attached to the same pole, we asked whether the decision to remove centromeric cohesin or not is taken already before entry into meiosis II. In other words, the chromosome itself is carrying the information of which “step” of step-wise cohesin removal (from arms or the centromere region) has to be executed. We found that bivalents segregating in meiosis I are getting prepared for sister separation in meiosis II. This preparation corresponds to the kinetochore individualization we describe here, taking place in anaphase I. By inhibiting arm cohesin removal through conditional knock-out of Separase, we obtained metaphase II oocytes harbouring bivalents that are attached in a monopolar manner, such as in metaphase I. This is not surprising, because bivalents are held together by chiasmata which lead to co-orientation of sister kinetochores and increase the likelihood of sister kinetochores being attached to the same pole (Herbert et al., 2015). Under these conditions the SAC in meiosis II is satisfied and oocytes can be activated, because even though bivalents and not dyads are present, they are attached and under tension. Crucially though, bivalents with fused kinetochores separate into chromosomes whereas bivalents with individualized kinetochores segregate into sister chromatids. Furthermore, the separated chromosomes behave like in meiosis I- they individualize kinetochores, resulting in dyads such as usually observed upon exit from meiosis I **(Fig. 7)**. Our data are in agreement with a recent study, showing that bivalents transferred from meiosis I into meiosis II oocytes behave as if they were in meiosis I and segregated into dyads (Ogushi et al., 2020).

**Figure 7.**
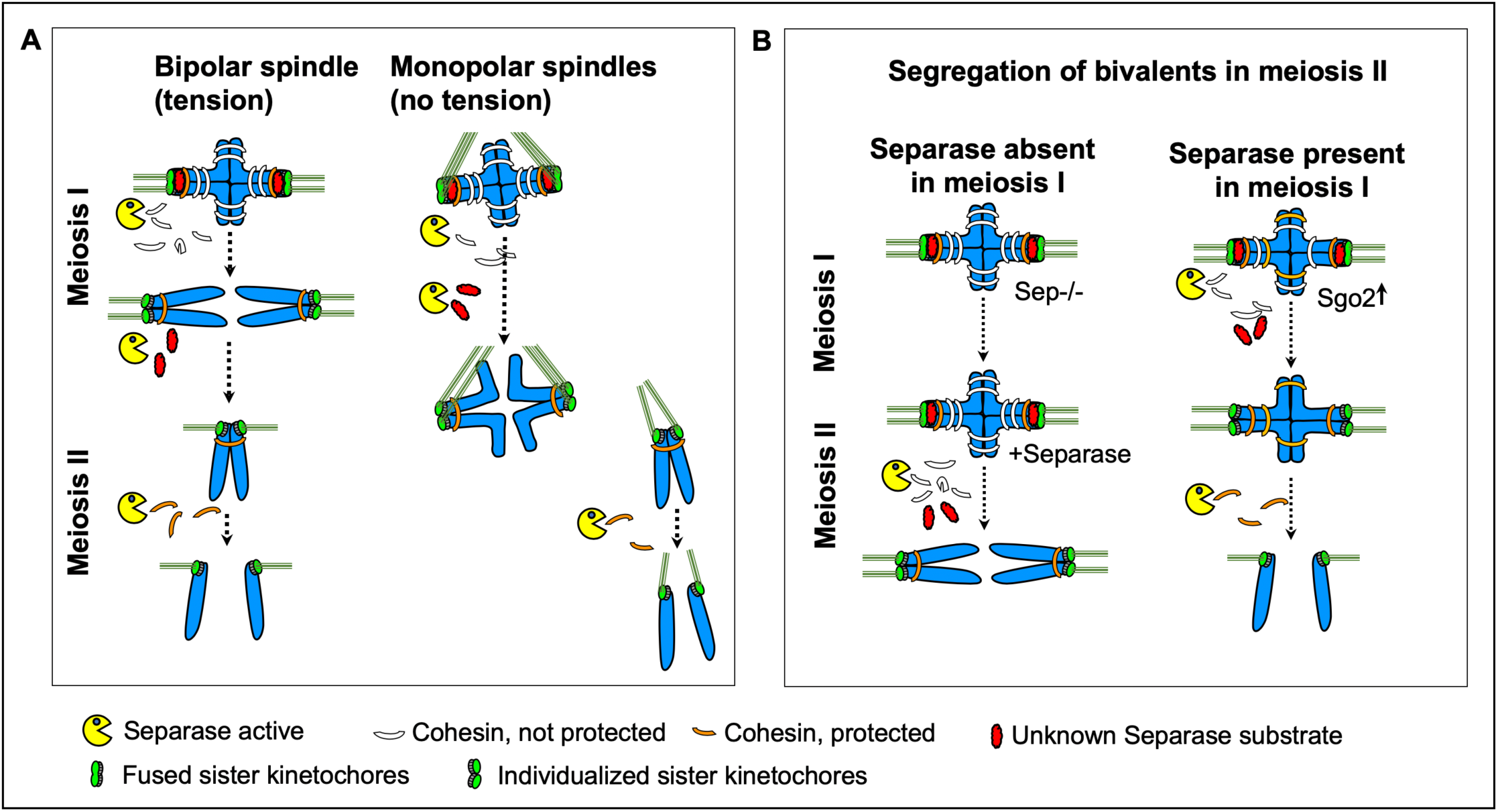
Kinetochore individualization in meiosis I is required for step-wise cohesin removal. On monopolar spindles, chromosomes segregate in meiosis I, and sister chromatids in meiosis II. **B)** Model of how Separase-dependent kinetochore individualization primes bivalents for deprotection of centromeric cohesin. If Separase is absent in meiosis I, arm cohesin and not centromeric cohesin is removed in meiosis II. Under these conditions, sister kinetochores individualize in meiosis II instead of meiosis I, demonstrating that step-wise cohesin removal (first arm cohesin, then centromeric cohesin) depends on the kinetochore structure. Without kinetochore individualization, centromeric cohesin cannot be cleaved in the subsequent division.

We found that kinetochore individualization depends on cleavage activity of Separase. What is the substrate of Separase that is holding kinetochores together until arm cohesin has been removed, usually in anaphase I? At this point we ignore its identity, but one attractive candidate is a separate fraction of Rec8, which is distinct from Rec8 on arms and Rec8 protected until anaphase II onset. Indeed, a third fraction of Rec8 has been proposed to exist and to confer loss of sister kinetochore co-orientation and meiosis II cohesin protection upon cleavage by Separase after meiosis I in a very recent study (Ogushi et al., 2020). According to our data, this unknown Separase substrate does not seem to be protected by Sgo2. How cleavage of this substrate is prohibited before anaphase I, when arm cohesin has been cleaved already, is therefore a mystery. Loss of Sgo2 in Sgo2 knock-out oocytes results in separation of sister chromatids instead of chromosomes in meiosis I (Llano et al., 2008), even though kinetochores should still be fused according to our hypothesis. Hence, kinetochore fusion cannot substitute for centromeric cohesin or centromeric cohesin protection, to prevent precocious sister chromatid segregation in meiosis I.

In conclusion, the decision to protect or deprotect centromeric cohesin is not due to the fact that oocytes are in meiosis I or II, nor mono-or bipolar attachment to the spindle, but lies within the chromosome itself. Importantly, we do not exclude that inactivation of Sgo2, or colocalization with I2PP2A/Set contribute to cleavage of centromeric Rec8 in meiosis II, but we propose that the key event that determines whether centromeric cohesin can be cleaved or not is sister kinetochore individualization. Unfortunately, these processes cannot be studied in mitotic cells, because even though Rec8 forms cohesive complexes when co-expressed with Stag3, these meiotic cohesin complexes are still removed by the prophase pathway and protected by Sgo1 (Wolf et al., 2018), which requires Bub1 kinase-dependent phosphorylation of Histone H2A, unlike Sgo2 (Marston and Wassmann, 2017). Hence also in the future, elaborate and technically challenging experiments in meiocytes will be required to gain further insights into the underlying molecular processes for cohesin protection and deprotection in meiosis.

## Material and Methods

### Animals

Mice were maintained under temperature, humidity and light controlled conditions under the authorization C75-05-13 at UMR7622 in a conventional mouse facility, with food and water access *ad libitum*. The project was submitted to ethical review according to the French law for animal experimentation (authorization B-75-1308). Adult CD-1 mice were purchased (Janvier, France) and C57BL/6 mice of the indicated genotypes were bred in our animal facility. Mice were not involved in any procedures except for genotyping prior to being sacrificed by cervical dislocation between 8-16 weeks of age, to dissect ovaries and harvest GV stage oocytes.

### Mouse oocyte culture

Oocytes were harvested in M2 Medium (Merck Millipore, MR-015P) supplied with 100µg/mL dibutyryl cyclic AMP (dbc-AMP; Sigma-Aldrich, D0260) to keep them arrested in GV stage until release through several wash steps into M2 medium without dbcAmp. Only oocytes undergoing GVBD up to 90 min after release were used. For microinjections, oocytes were kept 2-3 hours arrested before release. For longer incubations (more than three hours, e.g. Morpholino knock-down experiments) oocytes were put into M16 medium (Merck Millipore) containing dbcAmp, in CO_2_. All oocyte culture was done in medium drops covered with mineral oil (Sigma-Aldrich, M8410). To obtain oocytes in anaphase II, *in vitro* cultured metaphase II oocytes were artificially activated. Metaphase II oocytes were placed into home-made M16 medium without CaCl_2_ for 30min to 1h prior to being placed into the activation medium (M16 medium without CaCl_2_ supplied with 100mM Strontium chloride, Sigma-Aldrich 204463), for 1hour (except for protein detection during early anaphase II in Fig. 2E incubation time was reduced to 25min), in a CO_2_ incubator. To obtain *in vivo* matured oocytes in metaphase II, adult female mice were hormonally stimulated with a first injection of 3,3UI of pregnant mare serum gonadotropin (PMSG; Abbexa LTD, abx260389, followed by a second injection 48h later with 5UI of chorionic gonadotropin (HcG, Abbexa LTD, abx260092). 12h to 16h after the second injection, oocytes containing cumulus cells were collected from oviducts and put into M2 Medium. To collect oocytes, the medium was supplied with Hyaluronidase at 0,1 mg/mL (Sigma-Aldrich, H4272-30MG) for 5 to 10 min and oocytes were harvested by mouth pipetting.

### Drug treatment

All inhibitors were dissolved in DMSO (Sigma-Aldrich, D2650) and added to M2 or M16 medium (Merck Millipore, MR-10P). STLC (Sigma-Aldrich, 164739) was used at 1,5µM and 5µM for meiosis I and meiosis II experiments respectively, and Reversine (Cayman Chemical, 10004412) was used at 0,5µM and additionally added to the oil used to cover the culture droplets.

### In vitro transcription and microinjections

Microinjection pipettes were self-made, using a magnetic puller (Narishige; PN-31). Oocytes were manipulated on an inverted Nikon Eclipse Ti microscope, with a holding pipette (Eppendorf, 5178108.000), and injections were done using a FemtoJet Microinjector pump (Eppendorf) with continuous flow. mRNAs for injection into GV or CSF arrested metaphase II oocytes were transcribed using the mMessage mMachine T3 Kit (Invitrogen, AM1348) and purified using the RNeasy mini-kit (Qiagen, 74104). Plasmids to express Separase have been described (Kudo et al., 2006). GFP-Sgo2 was transcribed from the plasmid obtained by sub-cloning mouse Sgo2 (gift from K. Nasmyth) into pRN3- EGFP-C1 vector (Hached et al., 2011).

### Chromosome spreads and immunofluorescence

Oocytes were fixed at the indicated times after GVBD. To obtain metaphase II oocytes, oocytes were fixed 16h to 20h after GVBD, or after *in vivo* maturation, where indicated. Prior to oocyte harvesting, the *Zona pellucida* of oocytes was removed in successive baths in acidic Tyrode’s solution (home-made) without mineral oil, and oocytes were left to recover. For chromosome spreads, oocytes were fixed in 0,65% - 1% paraformaldehyde (Sigma-Aldrich, 441244), 0,15% Triton-X100 (Sigma Aldrich, T8787), and 3mM DTT (Sigma Aldrich, D9779) (Chambon et al., 2013a). For whole-mount staining of stable spindle microtubules oocytes were incubated 2-6 min in a cold treatment solution (80mM PIPES (Euromedex, 1124), 1mM MgCl_2_ (Euromedex, 2189-C) on top of an ice-water bath. They were immediately fixed for 30min in BRB80 buffer containing 0,3% Triton-X100 and 1,9% formaldehyde (Sigma-Aldrich, F1635). After several washes, oocytes were incubated over night at 4°C in a PBS-BSA 3% solution supplied with 0,1% Triton-X100 (Vallot et al., 2018).

The following primary antibodies were used at the indicated concentrations: human CREST serum auto- immune antibody (Immunovision, HCT-100, at 1:50), mouse monoclonal anti-PP2A C subunit clone 1D6 Alexa Fluor 488 conjugate (Sigma-Aldrich, 05-421-AF488, 1:50), polyclonal rabbit anti-Sgo2 antibody (gift from José Luis Barbero, 1:50), rabbit anti-REC8 (gift from Scott Keeney, 1:50), mouse monoclonal anti-α-tubulin (DM1A) coupled to FITC (Sigma-Aldrich, F2168, 1:100) polyclonal rabbit anti-Mps1 (gift from Hongtao Yu, 1:50).

Secondary antibodies were used at the following concentrations: donkey anti-human CY3 (709-166- 149, Jackson ImmunoResearch, 1:200), donkey anti-human Alexa Fluor 488 (709-546-149, Jackson ImmunoResearch, 1:200), donkey anti-mouse CY3 (715-166-151, Jackson ImmunoResearch, 1:200), donkey anti-rabbit CY3 (715-166-152, Jackson ImmunoResearch, 1:200), donkey anti-rabbit Alexa Fluor 488 (711-546-152, Jackson ImmunoResearch, 1:200), donkey anti-mouse Alexa Fluor 488 (715-546-150, Jackson ImmunoResearch, 1:200), donkey anti-human Alexa Fluor 647 (709-606-149, Jackson ImmunoResearch, 1:200), donkey anti-rabbit Alexa Fluor 647 (711-606-152, Jackson ImmunoResearch, 1:200), donkey anti-mouse Alexa Fluor 647 (715-606-150, Jackson ImmunoResearch, 1:200).

To stain chromosomes, Hoechst 33342 (Invitrogen, H21492) at 50µg/mL was used during secondary antibody incubation with AF1 Citifluor mounting medium (Biovalley, AF1-100), or IF Hardset + DAPI (VECTASHIELD) (Eurobio H-1200) to mount the slides.

### Image acquisition and Image treatment

To image chromosomes spreads, an inverted Zeiss Axiovert 200M microscope with a 100X/1,4 NA oil objective coupled to an EMCD camera was used. 6 z-sections with 0,4µm interval were taken. Live imaging was performed on the same microscope using a Plan-APO (63x/1.4 NA) oil objective (Zeiss). Prior to acquisition oocytes were preincubated in M2 media containing 1µM SirDNA (far-red DNA labeling probe, Spirochrome, SC007), for 1h. Oocytes in chambers were prepared for imaging as described previously (Nikalayevich et al., 2018). Time lapse was set up for 10 hours with 10 min intervals. 11 Z-stacks (3µm optical section spacing) were acquired in 491 nm channel for GFP and 640 nm channel for SirDNA and 1 z-section for the DIC image. To image whole-mount oocytes immunofluorescence, we used an inverted Leica laser scanning confocal microscope TCS SP5 II with a 63X oil immersion objective (HCX Plan APO CS, NA 1,4). Scan speed was 400Hz and Z-section interval was 0,08µm (Vallot et al., 2018). Pictures of cumulus were taken through a Zeiss Stereomicroscope Stemi2000 ocular and Asus ZenFone5Z camera (Sony IMX363 12MP). Image acquisitions were done with Metamorph or Leica Software. Images were processed using Fiji or ImageJ software (NIH). No manipulations were performed other than brightness and contrast adjustments, which were applied with the same settings to images that were compared.

### Quantifications and Statistical analysis

Quantifications were done using ImageJ. The mean fluorescence intensities were measured at centromeres on sum-projected images (by drawing a box around centromeres, as depicted on the scheme to each dot plot). The mean fluorescence intensities (I(fluor)) were corrected to background (bg) and normalized to CREST signals as follows: I(fluor)_normalized_ = [I(fluor)-I(fluor)^bg^] / [I(fluor)(CREST)- I(flour)^bg^(CREST)] Data plots were obtained with PRISM6 software. Quantification results were compared using the two-tailed Student’s t-test. All experiments were performed at least in experimentally independent duplicates. Shown are results from all replicates performed.

## Acknowledgements

We thank Marie-Hélène Verlhac and M-Emilie Terret for advice on oocyte culture, Camille Gauthier and Alison Kem-Seng for preliminary experiments, and members of the MOM group for advice and discussion. We thank Scott Keeney for Rec8 antibody, José Barberos for Sgo2 antibody, Jan van Deursen for the Bub1 kinase-dead mouse strain, Kim Nasmyth for the Separase conditional knock-out mouse strain and mouse Sgo2 plasmid, and Hongtao Yu for Mps1 antibody. L.K. was the recipient of a PhD fellowship by FRM (Fondation de la Recherche Medicale, ECO 20170637505). The KW lab obtained funding for this project by the ANR (ANR-16-CE92-0007-01, ANR-19-CE13-0015), FRM (Equipe FRM DEQ 20160334921), through “Emergence” funding by Paris Sorbonne Université (OoCyclins) and core funding by the CNRS and Sorbonne Université.

## Author contributions

Most experiments in Figures 1, 2, and 6 were performed by YG, experiments in Figure 3 by LK, and experiments in Figure 4 and 5 by SAT and DC. WEY and EB performed the experiment in Figure 2B and S1. DC provided expert technical help throughout the project. Figures were prepared by YG, LK, SAT and KW. Overall supervision, funding acquisition and project administration were done by KW and the manuscript was written by KW with substantial input from all authors.

## Figures

**Supplementary Figure 1.**
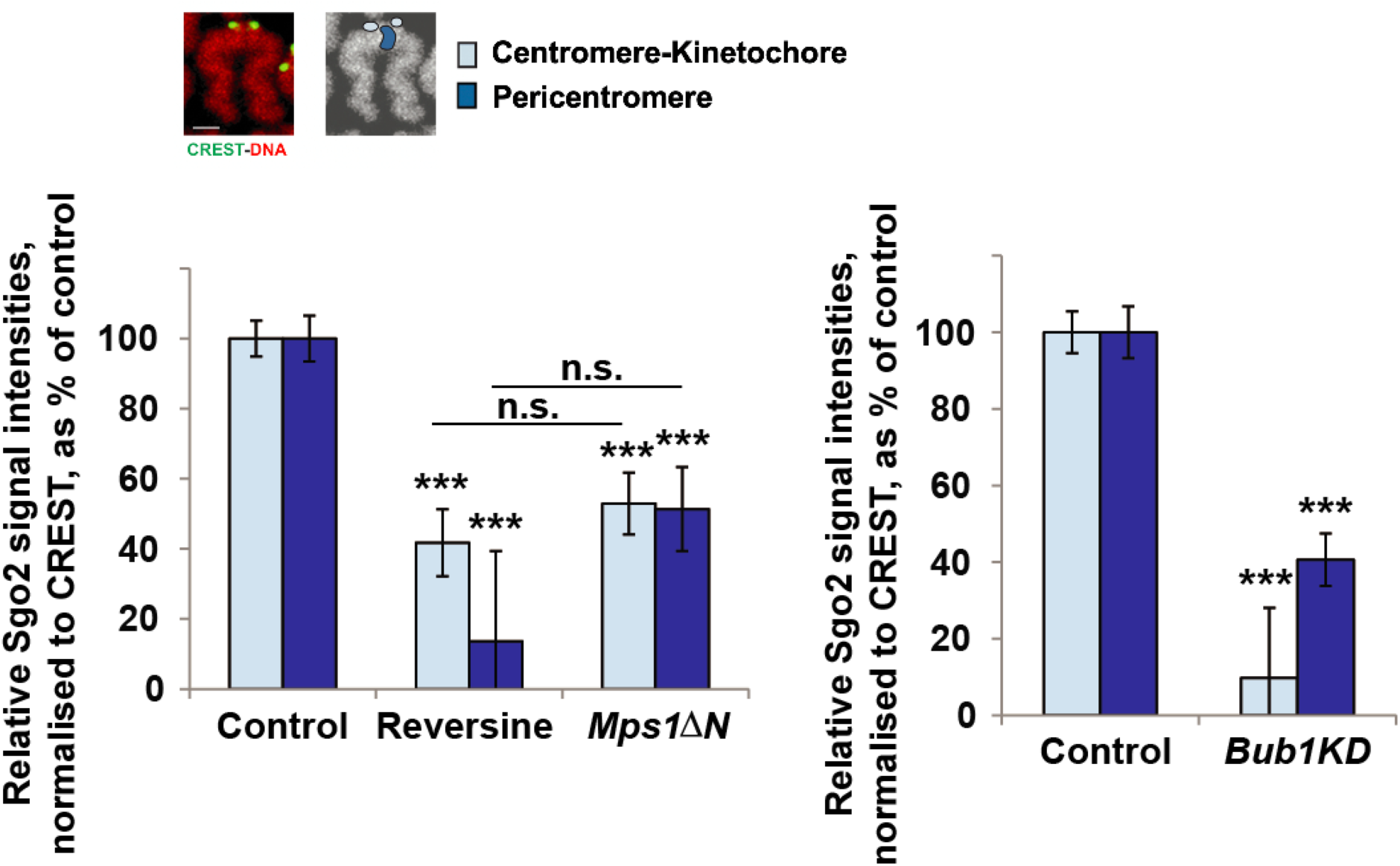
Quantification of the different pools of Sgo2 in meiosis II. The graphs show the corresponding quantification of the core centromere/ kinetochore signal (light blue: overlapping with CREST) and the pericentromere signal (dark blue: not overlapping with CREST) for Figure 2B. Mean of Sgo2/CREST intensities at the centromere versus pericentromere in *Bub1KD*, Reversine-treated and *Mps1ΔN* oocytes were normalised to the mean Sgo2/CREST intensities in control oocytes. Number of kinetochore pairs and number of oocytes analysed are indicated. Values indicate mean, error bars ± s.e.m. from one representative experiment performed at least in duplicate, using Student’s *t*-test. (n.s.: not significant; \*\*\**P*<0.0001).

**Supplementary Figure 2.**
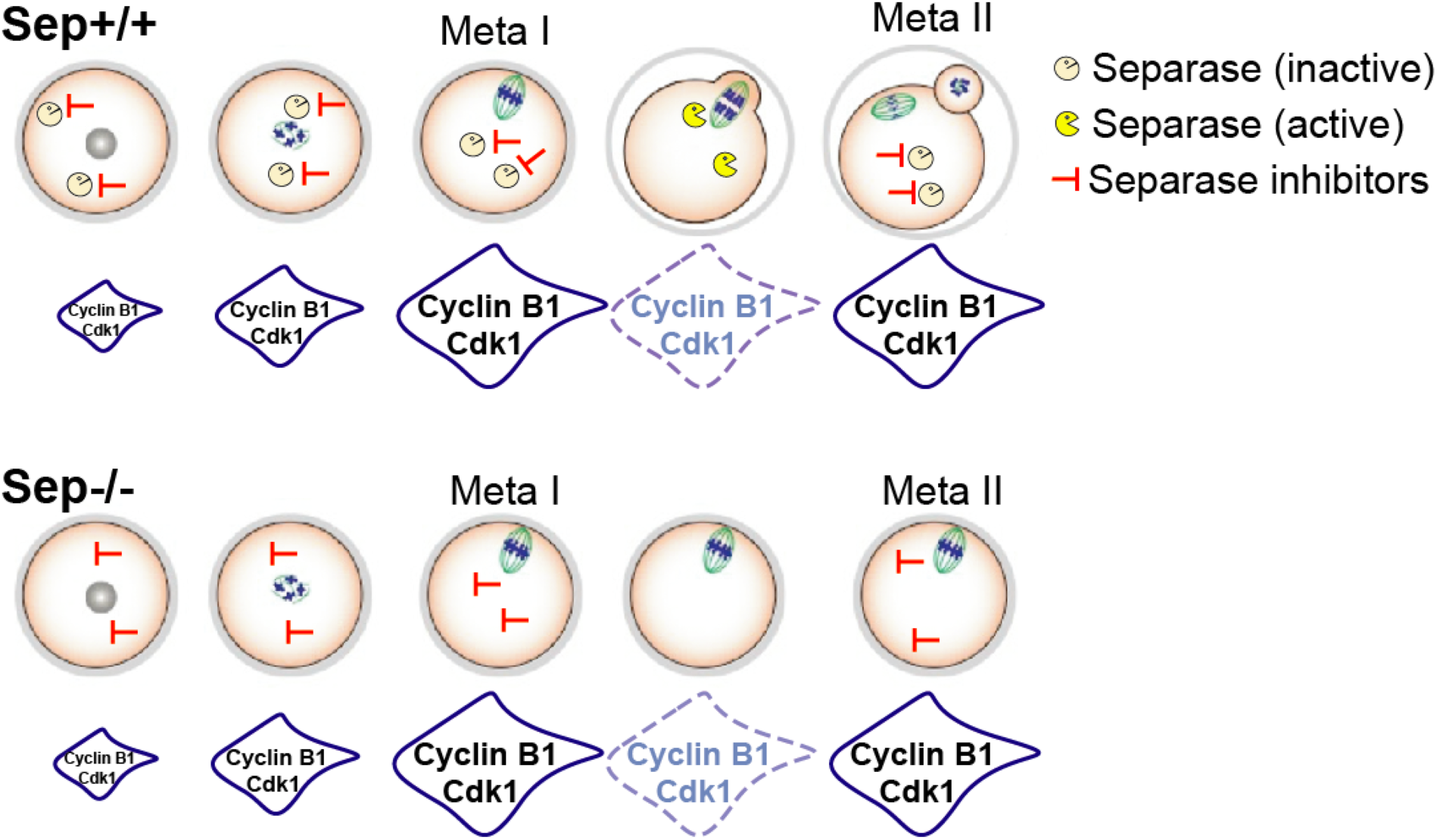
Oocytes harbouring a conditional knock-out of Separase progress into meiosis II. Scheme summarizing published data on cell cycle progression in oocytes devoid of Separase. Cdk1 activity accumulates, disappears, and re-accumulates on time. The APC/C substrate Cyclin B1 is degraded on time in the absence of Separase, and re-accumulates as oocytes progress into meiosis II, even though no polar body extrusion is observed, due to a role of Separase in Polar body extrusion, which is independent of its cleavage activity. The spindle assembly checkpoint is not activated, because chromosomes are correctly attached without Separase. See text for references.

**Supplementary Figure 3.**
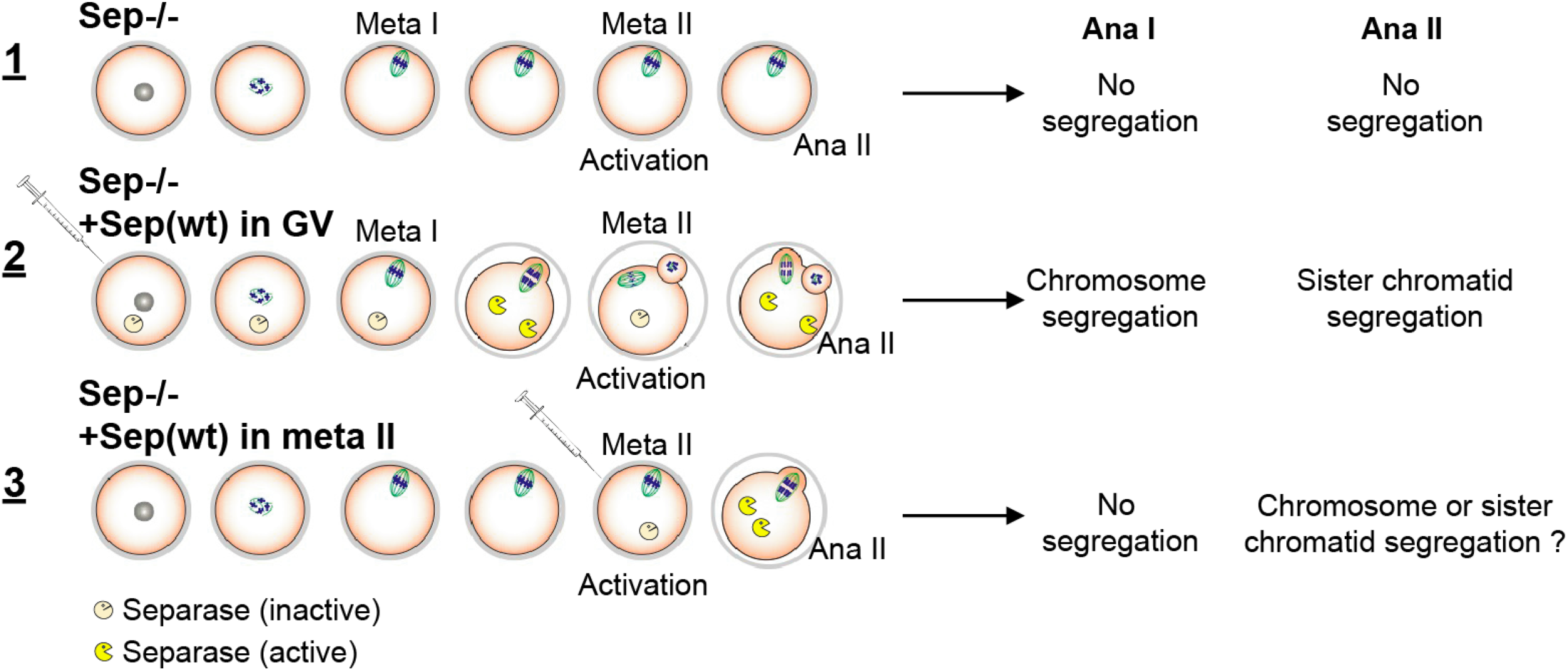
Scheme of rescue experiment in Sep-/- oocytes in meiosis I or II. 1) Sep -/- oocytes without rescue. 2) Sep-/- oocytes rescued from GV onwards (Figure 4). 3) Sep-/- oocytes rescued from metaphase II onwards. The question we asked was whether chromosomes or sister chromatids are separated when Separase is absent in meiosis I, and present in meiosis II.

